# Individual level predictions of *Staphylococcus aureus* bacteraemia-associated mortality

**DOI:** 10.1101/071837

**Authors:** Mario Recker, Maisem Laabei, Michelle S. Toleman, Sandra Reuter, Beth Blane, Estee Török, Sion Bayliss, Sharon J. Peacock, Ruth C. Massey

## Abstract

The bacterium *Staphylococcus aureus* is a major human pathogen, where the emergence of antibiotic resistance is a global public-health concern. Host factors such as age and the presence of co-morbidities have been implicated in a worse outcome for patient. However, this is complicated by the highly complex and multi-faceted nature of bacterial virulence, which has so far prevented a robust mapping between genotype, phenotype and infection outcome. To investigate the role of bacterial and host factors in contributing to *S. aureus* bacteraemia-associated mortality we sequenced a collection of clinical isolates (of the MLST clonal complex CC22) from patients with bloodstream infections and quantified specific virulence phenotypes. A genome-wide association scan identified several novel virulence-affecting loci, which we validated using a functional genomics approach. Analysing the data comprising bacterial genotype and phenotype as well as clinical meta-data within a machine-learning framework revealed that mortality associated with CC22 bacteraemia is not only influenced by the interactions between host and bacterial factors but can also be predicted at the individual patient-level to a high degree of accuracy. This study clearly demonstrates the potential of using a combined genomics and data analytic approach to enhance our understanding of bacterial pathogenesis. Considering both host and microbial factors, including whole genome sequence and phenotype data, within a predictive framework could thus pave the way towards personalised medicine and infectious disease management.

## Introduction

*Staphylococcus aureus* bacteraemia (SAB) is a significant global health problem^1^ and is exacerbated by the emergence and widespread circulation of drug resistant strains, such as methicillin-resistant *S. aureus*, or MRSA^2^. Mandatory surveillance of SAB has now been implemented in several countries, with many reporting a decline in the incidence of methicillin-resistant SAB (MR-SAB)^3–5^. However, in the UK the incidence of methicillin-susceptible SAB (MS-SAB) has been increasing year on year, where there has been a 15.4% increase in cases since reporting became mandatory in 2011/2012^3^. Furthermore, the 30 day (all-cause) mortality rate for SAB has not significantly changed over the last two decades and appears to have plateaued at approximately 20%^6^. This strongly suggests that existing infection control and treatment options are insufficient to tackle this important health problem and that a better understanding of the factors that contribute to bacteraemia-associated morbidity is crucially needed.

To date, many host risk factors have been identified for both the occurrence of and treatment failure following SAB^6^. However, the contribution of the bacterium is only partially understood and is largely informed by experimental animal studies. These model systems come with their own set of limitations, and many observations from these contrast with those from human studies. For example, whereas cytolytic toxins have previously been shown to enhance disease severity in animal models of SAB^7,8^, isolates from invasive diseases in humans, such as bacteremia and pneumonia, were recently found to be significantly less toxic than those isolated from skin and soft tissue infections or even those of healthy volunteers^9–12^. This raises the question as to whether animal models are adequate to study bacterial virulence in human SAB infections, or whether there is an important distinction between the role of toxicity in causing bacteremia and its pathogenic effect once bacteremia has been established. Either way, human-based approaches are essential to close this gap in our knowledge.

Another limitation to our understanding of the pathogenesis of SAB is that most studies focus on only on a single or small number of factors in isolation, host or bacterial. For example, several studies have found increased mortality rates associated with MR-SAB compared to MS-SAB^13,14^. However, patients with co-morbidities are more likely to develop an MR-SAB due to their impaired health and longer time spent in healthcare facilities when compared to those without comorbidities. When subsequent studies considered both factors together, no difference in mortality was associated with the methicillin resistance status of the infecting bacterium^15^. This illustrates the importance of a more inclusive approach that considers all of the potential host and bacterial factors and examines how their interactions influence the outcome for the patients.

We have previously demonstrated how genotype-phenotype mapping in *S. aureus* has the potential to provide sufficient information to enable predictions of the level of virulence expressed by the infecting microorganism^16^. Here, by expanding this whole-genome approach to a set of fully sequenced isolates from bacteraemic patients and analysing the combined set of genotype, phenotype and clinical meta-data within a machine learning predictive modelling framework we show how host and bacterial factors interact to determine severe infection outcome of *S. aureus* bacteraemia. These findings pave the way towards individual patient-level predictions and personalised treatment strategies.

## Material and Methods

### Strain and clinical metadata collection

All isolates were collected from adults admitted to a single hospital with their first episode of SAB between 2006 and 2012, and were stored in glycerol at −80°C. Samples were collected during an observational cohort study of adults with SAB at Addenbrooke’s Hospital, Cambridge, UK between 2006 and 2012. Written informed consent was not required as the study was conducted as part of a service evaluation of the management of SAB. Ethical approval was obtained from the University of Cambridge Human Biology Research Ethics Committee (reference HBREC.2013.05) and the Cambridge University Hospitals NHS Foundation Trust Research and Development Department (reference A092869). Study definitions have been defined previously^17^ and were used to determine the focus of the bacteremia, classify the bacteremia as community-acquired, hospital-acquired or healthcare-associated, and to report outcomes, including death at 30 days. The presence of comorbidities was assessed using the Charlson comorbidity index (CCI) and were dichotomized into scores of <3 or ≥3 ^18,19^, as detailed in Supplementary Table 1.

### Genome Sequencing

Bacterial DNA extraction was carried out on a QIAxtractor (Qiagen), and library preparation was performed as previously described^20^. Index-tagged libraries were created, and 96 separated libraries were sequenced in each of eight channels using the Illumina HiSeq platform (Illumina) to generate 100-bp paired-end reads at the Wellcome Trust Sanger Institute, UK. Paired-end reads for these isolates were mapped to the ST22/EMRSA15 reference strain, HO 5096 0412^21^ and SNPs were identified as described previously^21^. The accession number for the sequence data for each of these isolates is listed in Supplementary Table 1.

### Cytotoxicity Assay

Overnight *S. aureus* cultures were diluted 1:1000 into fresh tryptone soya broth and incubated for 18 h at 37°C with shaking at 180 rpm in 30 mL glass tubes. *S. aureus* supernatants were harvested from 18 h cultures by centrifugation at 14,600 rpm for 10 min. The THP-1 human monocyte-macrophage cell line (ATCC#TIB-202) was routinely grown in suspension in 30 mL of of RPMI-1640, supplemented with 10% heat-inactivated fetal bovine serum (FBS), 1 μM L-glutamine, 200 units/mL penicillin, and 0.1 mg/mL streptomycin at 37°C in a humidified incubator with 5% CO_2_. Cells were routinely viewed microscopically every 48–60 h and harvested by centrifugation at 1,000 rpm for 10 min at room temperature and re-suspended to a final density of 1–1.2 x 10^6^ cells/mL in tissue-grade phosphate buffered saline. This procedure typically yielded >95% viability of cells as determined by trypan blue exclusion and easyCyte flow cytometry. To evaluate *S. aureus* toxicity, we diluted the supernatant to 30% of the original volume in TSB and incubated 20 µL of washed THP-1 cells with 20 µL diluted bacterial supernatant for 12 min at 37°C under static conditions. Cell death was quantified by easyCyte (Millipore) flow cytometry using the Guava Viability stain (Millipore) according to manufacturer’s instructions, with the toxicity of each isolate quantified in triplicate and the mean of this presented (listed in Supplementary Table 1).

### Biofilm assay

Biofilm formation was quantified using a 1:40 dilution from overnight cultures into 100 µL of fresh tryptic soy broth supplemented with 0.5% sterile filtered glucose (TSBG) in 96-well polystyrene plate (Costar). Perimeter wells of the 96-well plate were filled with sterile H_2_O and plates were placed in a separate plastic container inside a 37°C incubator and grown for 24 h under static conditions. For the transposon mutants, erythromycin (5 µg/mL) was added to the growth medium. Semi-quantitative measurements of biofilm formation on 96-well polystyrene plates was determined based on the method of Ziebuhr et al^22^. Following 24-h growth, plates were washed vigorously five times in PBS, dried and stained with 150 μL of 1% crystal violet for 30 min at room temperature. Following five washes of PBS, wells were re-suspended in 200 μL of 7% acetic acid, and optical density at 595 nm was recorded using a Fluorimeter plate reader (BMG Labtech). To control for day to day variability, for the clinical isolates a control strain (E-MRSA15) was included on each plate in triplicate, and absorbance values were normalised against this (listed in Additional Table 1). For the transposon mutants, as JE2 is the wild type strain this was used as the control strain, and the effect of mutating the loci made relative to this. For this experiment the assays were performed in triplicate on each plate and repeated four times.

### Genome wide association study (GWAS)

The genome of the reference strain HO 5096 0412 was split into 2095 variable loci, corresponding to coding region and intergenic regions, containing SNPs relative to the reference genome. Annotated intergenic elements such as miscellaneous RNAs were considered separate loci. Synonymous SNPs in coding regions and SNPs in known mobile genetic elements and repeat regions were not considered. The resulting loci were named allele_X where X refers to the position of the SNP at the 5’ end of that block, relative to the origin of replication. Each of these alleles in each isolate was scored as 1 if it differed from the reference and 0 if it didn’t. These allele scores for each isolate were used as the genotypic information for the following analysis. Significant associations between bacterial genotype and either phenotype (toxicity and biofilm formation) were identified by fitting an analysis of variance model (ANOVA) in R^23^ and using a minor allele frequency cut-off of 5%. The reported *P* values are not corrected for multiple testing; the Bonferroni statistical significance threshold is instead provided in fig. 2.

### Predictive modelling

We employed a *Random Forests*^24^ machine learning approach, using the *randomForest* package in R^25^, to identify predictive signatures of host mortality based on the genotype, phenotype and clinical meta-data. To assess the models’ predictive accuracies we employed two different measures: (i) the receiver operating characteristic (ROC) curve, which is generated by plotting the true positive rate against the false positive rate (i.e. the observed incidence against the false predicted incidence) at various threshold settings and where the area under the curve (AUC) is a measure of predictive accuracy, with an AUC=1 equating to zero error and an AUC=0.5 equating to random guessing; and (ii) by means of a confusion matrix, which contrasts the instances of the predicted classes (*alive* or *death*) against the actually observed classes. The misclassification-rates reported here are based on the so-called out-of-bag errors^25^, which are derived by iteratively testing the models’ performances against subsets of data left out during the fitting processes and thus provide a measure of how well the models would fare against unknown data.

## Results

To elucidate the role of bacterial factors in human SAB we analysed a collection of 135 *S. aureus* isolates sampled from patients with bloodstream infections admitted to a single hospital between 2006 and 2012. All isolates belonged to the multi-locus defined clonal complex 22 (CC22) and contained both methicillin-resistant (MRSA) and methicillin-susceptible (MSSA) isolates. The 30-day all-cause mortality rate was 24.1% and the clinical data are summarized in Supplementary Table 1. Each isolate was whole-genome sequenced, and quantitatively phenotyped with respect to cytolytic activity (the ability to lyse a monocyte cell line, THP-1) and biofilm formation, both of which are major virulence determinants implicated in *S. aureus* disease^1,2^. Cytolytic toxins enable the evasion of cellular aspects of host immunity, release nutrient from host cells and are responsible for much of the purulent tissue damage associated of *S. aureus* infections^1,2^. Biofilm formation enables *S. aureus* to colonise foreign material and medical devices, protects the bacteria from many aspects of host immunity and renders some antibiotics less effective^1,2^. Despite the close genetic and geographic relationship between these isolates, toxicity and biofilm formation varied widely across the isolates; no association was found between methicillin susceptibility and either virulence phenotype (fig. 1).

**Figure 1.**
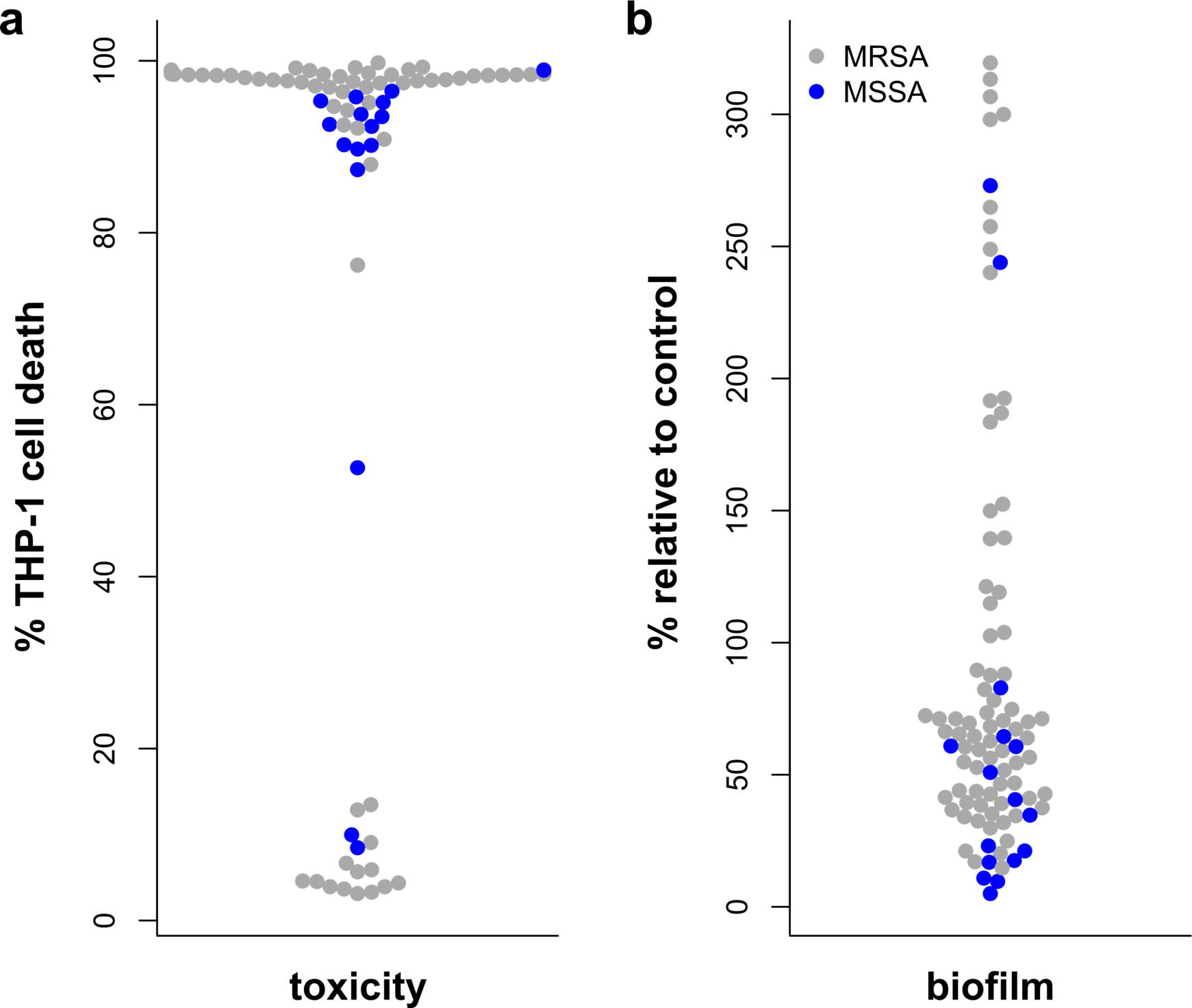
The toxicity and biofilm forming abilities of 135 *S. aureus* bacteraemia isolates. **(a)** Toxicity for each isolate was determined by incubating bacterial supernatant with cultured human cells, using flow cytotometry to quantify cell death (toxicity). No difference was observed between methicillin-resistant (MRSA, grey circles) and methicillin-susceptible (MSSA, blue circles) isolates. **(b)** Biofilm forming abilities were quantified relative to a control included in each assay in a static 96 well format. A wide range of biofilm forming abilities was evident with no discernible difference between methicillin-resistant (MRSA, grey circles) and methicillin-susceptible (MSSA, blue circles) isolates.

We performed a genome-wide association study (GWAS) to identify loci that were associated with the toxicity of the isolates and their ability to form biofilm, where associations were tested at an uncorrected (P<0.05) and a Bonferroni corrected (P<4.6x10^−5^) significance threshold (fig. 2a). For toxicity, no loci reached significance when using the Bonferroni correction for multiple comparisons, although mutations in the Agr toxicity-regulating locus and a putative membrane protein (SAEMRSA15_10750) were marginally associated (0.05 > *P* > 4.6x10^−5^). For biofilm, only one locus reached statistical significance, *pbp2*, a penicillin binding protein. This gene is believed to be essential to the bacteria, as no transposon mutants were available to functionally verify its effect on biofilm. However, members of this family of proteins have been shown previously to affect biofilm formation^26^. A further 37 loci showed marginally significant associations with biofilm (Supplementary Table 2), of which we chose a subset of 12 for functional validation using transposon mutants^27^ (Fig. 2b). Out of these, transposon mutagenesis of six loci showed a significant effect on biofilm formation compared to the wild type: a putative helicase (SAEMRSA15_23880, *Tn* mutant NE513), the quinolone efflux protein NorA (SAEMRSA15_16820, Tn mutant NE1034), a putative thiamine pyrophosphate enzyme/indole-3-pyruvate decarboxylase (SAEMRSA15_01530, *Tn* mutant NE1149), and a putative peptidase (SAEMRSA15_04760, *Tn* mutant NE1455), a putative inosine-uridine preferring nucleoside hydrolase (SAEMRSA15_02020, *Tn* mutant NE1637, and a hypothetical protein (SAEMRSA15_11850, *Tn* mutant NE318) (fig. 2b).

**Figure 2.**
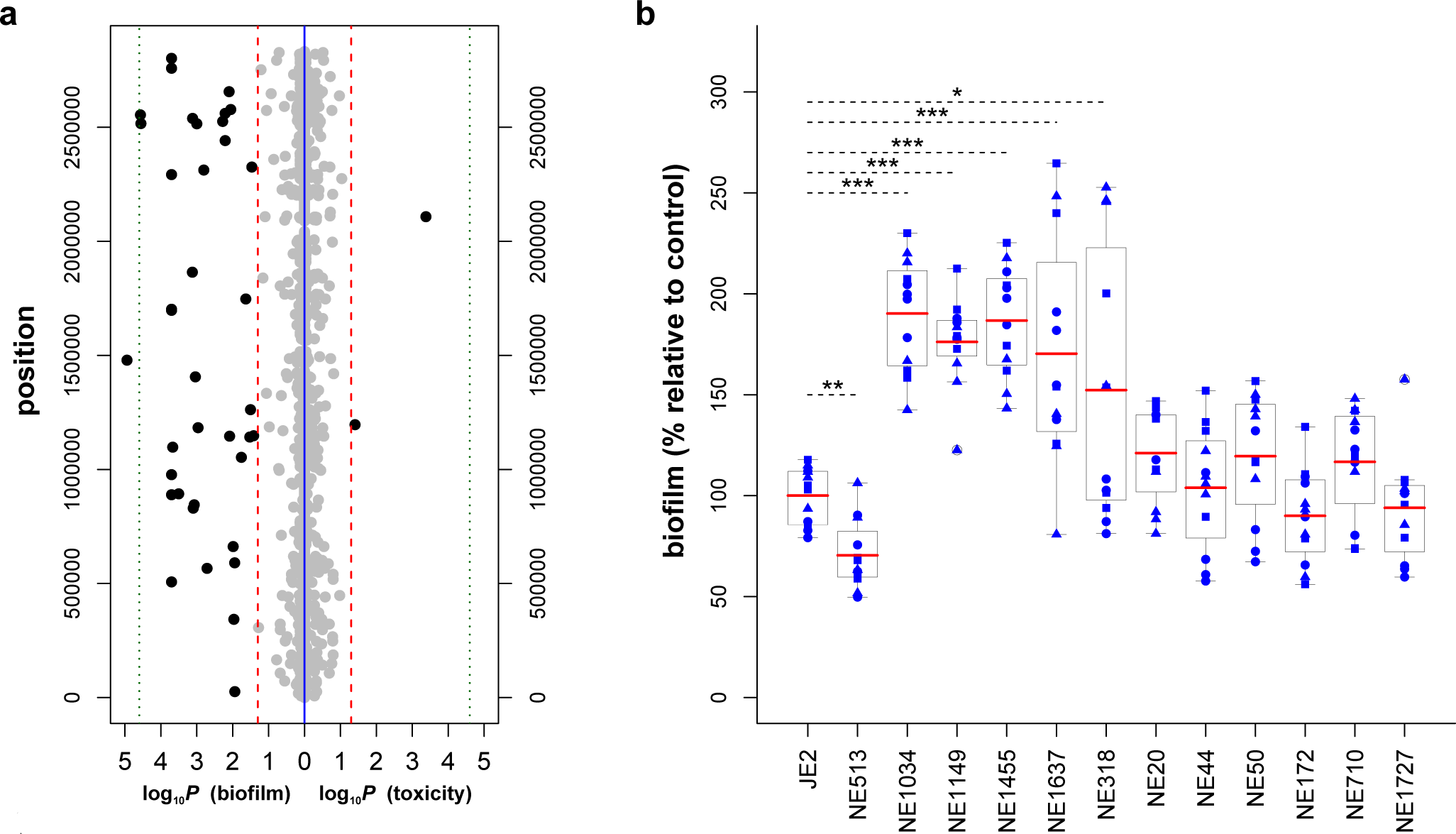
Genome-wide associations and functional validation of biofilm affecting polymorphisms. **(a)**Manhattan plots representing the results of the GWAS for both biofilm (left-hand side) and toxicity (right-hand side), performed on the 135 bacteremia isolates. For toxicity, only two polymorphic loci were significantly associated using an uncorrected threshold (indicated by the red vertical dashed line), one in the *agrC* gene and the other in a gene encoding a putative membrane protein. No loci were significantly associated with toxicity when Bonferroni was used to correct for multiple comparisons (indicated by the green vertical dotted line). For biofilm, one locus, the *pbp2* gene, was significantly associated using the the Bonferroni threshold, and a further 36 using the uncorrected threshold. **(b)** Twelve of the putative biofilm-affecting loci (associated by GWAS) were functionally validated using transposon insertion, of which six showed a significant effect compared to the wild-type (JE2) (Welch’s two-sided *t*-test, with *: *P* <2E-2, **: *P* <2E-4, ***: *P* <2E-6). Results are based on four biological replicates per strain that were repeated three times each.

Having access to the genetic and phenotypic data for each isolate as well as clinical data (Supplementary Table 1) for 92 of the 135 patients, we sought to determine the factors most predictive of severe infection outcome, namely host mortality at day 30. Due to the high dimensionality and high number of possible interactions between the various host and bacterial factors we employed a *random forests* machine learning approach^24,25^. We compared four scenarios, where the model was fitted against (i) the bacterial genotype data only, (ii) the bacterial phenotype data only, (iii) the clinical data only, and (iv) the entire dataset. Where genotype data were incorporated we first performed a feature selection process, in which the model was fitted against the entire dataset, and a smaller subset of variables selected for further analysis based on their predictive importance, i.e. their contribution to the model’s performance in distinguishing between the two classes. This reduced the total number of variables used for fitting down to just 20 (from over 2000 initially).

As shown by receiver operator characteristic (ROC) curves in figure 3a, data comprising either just bacterial factors (SNP or phenotype) or just clinical factors were relatively poor predictors of mortality. The most accurate model was one that included a combination of all factors, suggesting that these may interact in determining infection outcome. In this case, the model yielded a remarkable individual patient-level predictive accuracy of nearly 80% (based on out-of-bag error rates (which is a measure of prediction error)). This is shown by the illustrated confusion matrix in fig. 3b, where the dark-blue diagonal sectors represent the true positive and true negative rates and the light-blue off-diagonal entries represent the false positive and false negative rates.

**Figure 3.**
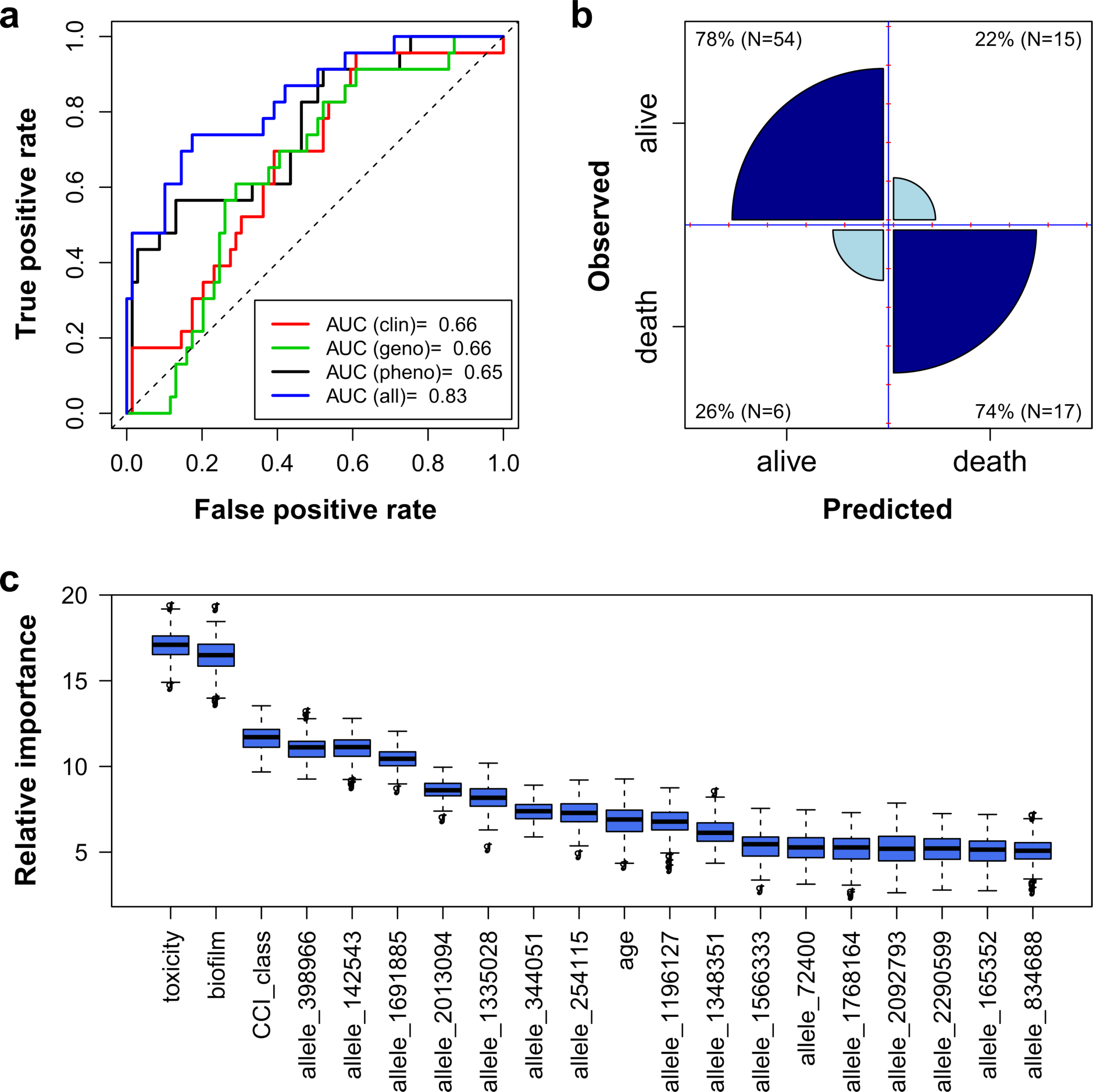
Predictive model performance and variable importance. **(a)** Receiver operator characteristic (ROC) curve of the *random forests* fit to four sets of variables consisting of clinical metadata (red line), genotype data (green line), phenotype data (black line) and all available variables (blue lie). As indicated by the area under curve (AUC), the model fit to a combination of all available data showed the highest predictive accuracy. **(b)** Confusion matrix illustrating the accuracy of the model in predicting individual patients 30-day mortality. The out-of-bag classification and misclassification rates (dark-blue/diagonal and light-blue/off-diagonal wedges, respectively) of the *random forests* model fitted to a feature-selected subset of the combined genotype, phenotype and clinical data. **(c)** Relative importance of the 20 features used for the final model, as measured by the random forest by means of a variable’s influence on the model’s predictive performance and based on 200 model fits, clearly demonstrating the significant effect of the bacteria’s phenotype on SAB-associated mortality.

*Random forest* can also be used to rank factors based on their relative importance to the predictive power of the model. As illustrated in fig. 3c toxicity and biofilm were the most important features, with patient age and co-morbidities (CCI-class) also playing important roles. Our model also identified a number of other bacterial genetic loci as contributing to disease outcome (listed in Supplementary Tables 3). Of particular note is the *capA* gene (indicated as allele_142543 in fig. 3c), which encodes a capsule biosynthesis enzyme. While the role of capsule in protecting bacteria against several aspects of the host immune system during experimental infections has been established^28,29^, several clinically important clones have evolved to be capsule-negative, due to polymorphisms in the capsule biosynthesis genes^30^. The protective activity of capsule is therefore not critical to the ability of *S. aureus* to cause disease in humans. Although we have not functionally verified the effect of the polymorphism found here on capsule expression, it suggests that this bacterial phenotype can also vary within a clone and potentially affect infection outcome.

Based on the ranked importance scores of the different factors used in this model (fig. 3c), both toxicity and biofilm formation are highly influential in determining SAB-associated mortality, the former positively and the latter negatively. To further analyse these and illustrate their influence alongside the other important factors we used the *random forest* approach again to derive two-way partial-dependence plots. As shown in figure 4, the combination of age with high levels of toxicity and/or low levels of biofilm was associated with a significant increase in the risk of patient death. Equally, the comorbidity index (CCI)^18,19^ with any combination of old age, high toxicity, and low levels of biofilm formation considerably increased the probability of severe infection outcomes.

**Figure 4.**
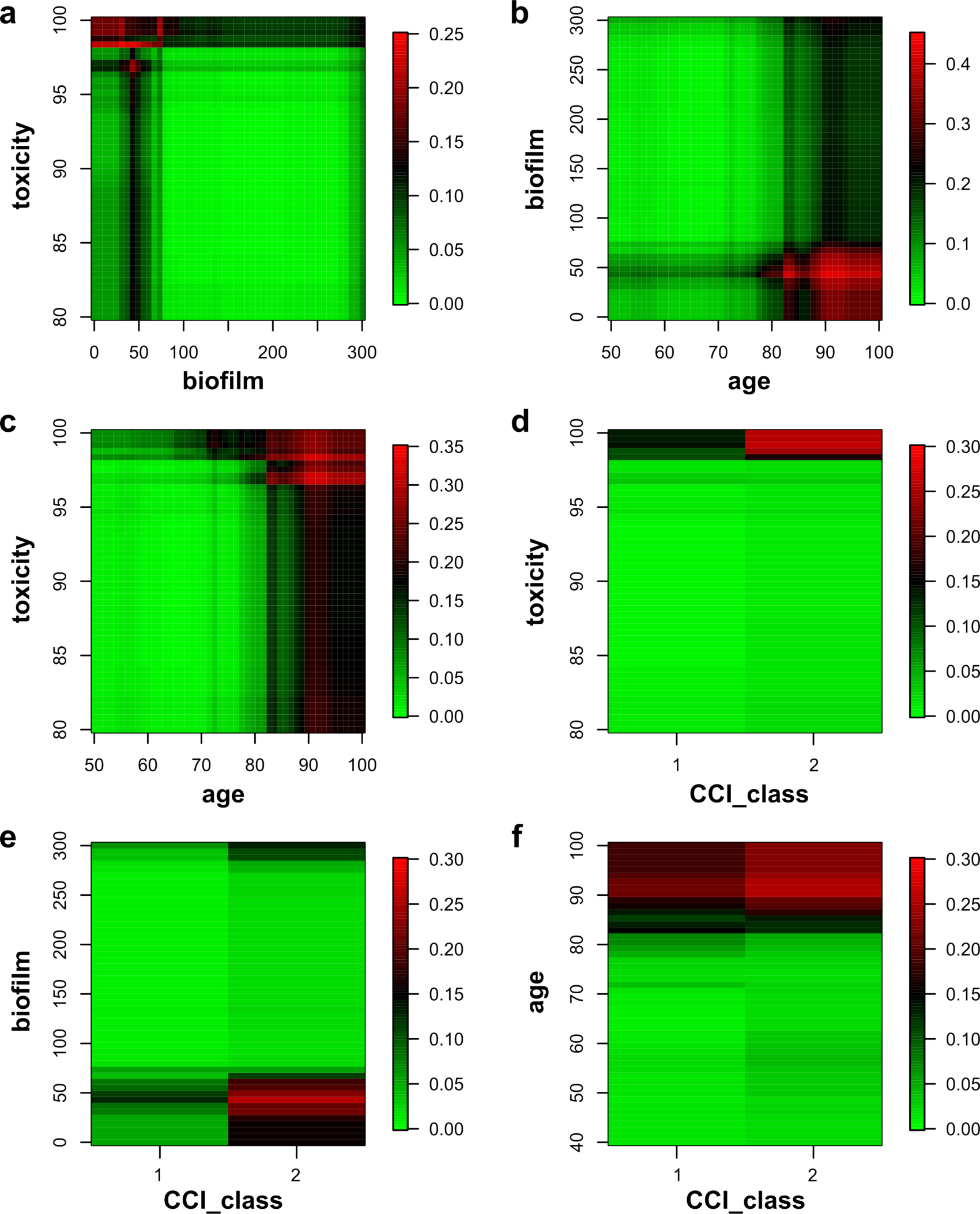
Interactions of phenotype and host factors determine risk of mortality. Two-way interaction plots derived by the *random forests* visualising the interactions between the two phenotype measures (toxicity and biofilm) and two important host factors and their combined effect on host mortality. The color bars indicate the risk of mortality, keeping all other variables fixed at either their means (for continuous variable) or their most common value (for categorical predictors).

## Discussion

In this study we demonstrate how combining genotypic, phenotypic and clinical metadata within a mathematical framework can be used to accurately predict the mortality of individual patients following SAB. This work also highlighting the critical role that specific bacterial phenotypes play in affecting the outcome from SAB, providing clarity to an apparent contradiction between experimental animal and human studies. Animal models have demonstrated a clear role for toxins in the severity and disease outcome from SAB^7,8^, however work by us and several other groups have found a negative correlation between toxicity and invasive diseases such as SAB in humans^9–12^. Here our data appears to conflict with our previous work and support the findings of the animal-based studies, which we believe can be reconciled by proposing a sequential pathway of events that distinguishes between the establishment of SAB (i.e. gaining access to the bloodstream) and the subsequent processes associated with a fatal outcome once SAB has become established. Toxin production appears to play disparate roles in these two stages of the disease, being selected against for the initial process of gaining access to the blood stream, but once in the bloodstream it is a major factor leading to host mortality, likely due to the damage and inflammation it causes there. A likely explanation for why animal models of SAB have only confirmed the latter process (disease severity) is that the natural infectious process of gaining access to the bloodstream is bypassed by injecting high doses (typically 10^7^-10^8^ colony forming units) directly into the animal’s bloodstream.

From the host perspective, the importance of patient age and co-morbidities in contributing to mortality is consistent with previous findings. However, our modeling approach also demonstrates that these host features can critically interact with bacterial factors and, importantly, allow a prediction of patient outcome with a high degree of accuracy. It is likely that other factors not considered here may influence a patient’s risk of death following SAB. Apart from patient care, host genotype and other bacterial phenotypes could have a significant effect on infection outcome. A larger more detailed dataset may enable us to fully identify these factors and unravel their interactions to predict mortality with even higher accuracy. However, given the known epidemiology (with older patients with co-morbidities being the most susceptible) and the complexity of this type of disease, it is unlikely that we will achieve 100% predictability, demonstrating the significance of the near 80% accuracy reported here.

With the growing global problem of antimicrobial resistance, alternative intervention and control strategies are needed. These include the development of vaccines and identification of drugs that attenuate the virulence of pathogens. However, without a full understanding of how the bacterial targets for these strategies are involved in causing disease in humans, there is a significant risk of investing in and pursuing unsuccessful lines of therapeutic development. Our findings here, for example, suggest that cytolytic toxins, components of biofilm and possibly capsule are unlikely to be good targets, as they play disparate roles in different stages of disease and their expression is highly variable, even within closely related bacterial clones. Of greater importance, however, it that this work has the potential to make a significant contribution to infectious disease diagnosis and management. With the move towards the introduction of microbial genome sequencing into routine diagnostic settings, the ability to use such information to inform clinicians on the likely outcome of an infection for individual patients would allow them to tailor treatment to that individual, an important step towards the implementation of personalised medicine approaches to infectious disease management.

## Acknowledgements

We would like to thank Dr Rebecca Saunderson, for her role in collecting and supplying the clinical data, and Dr Theodore Gouliouris, Dr Emma Nickerson and Dr Sani Aliyu for their role in collection of clinical information. We thank the Sanger Institute’s core Pathogen Production Groups and Pathogen Informatics Group.

## Supporting Information Captions

**Supplementary Table 1:** Strain information including clinical metadata, genome accession codes, toxicity and biofilm formation. (this has been provided as an excel spreadsheet)

**Supplementary Table 2:**
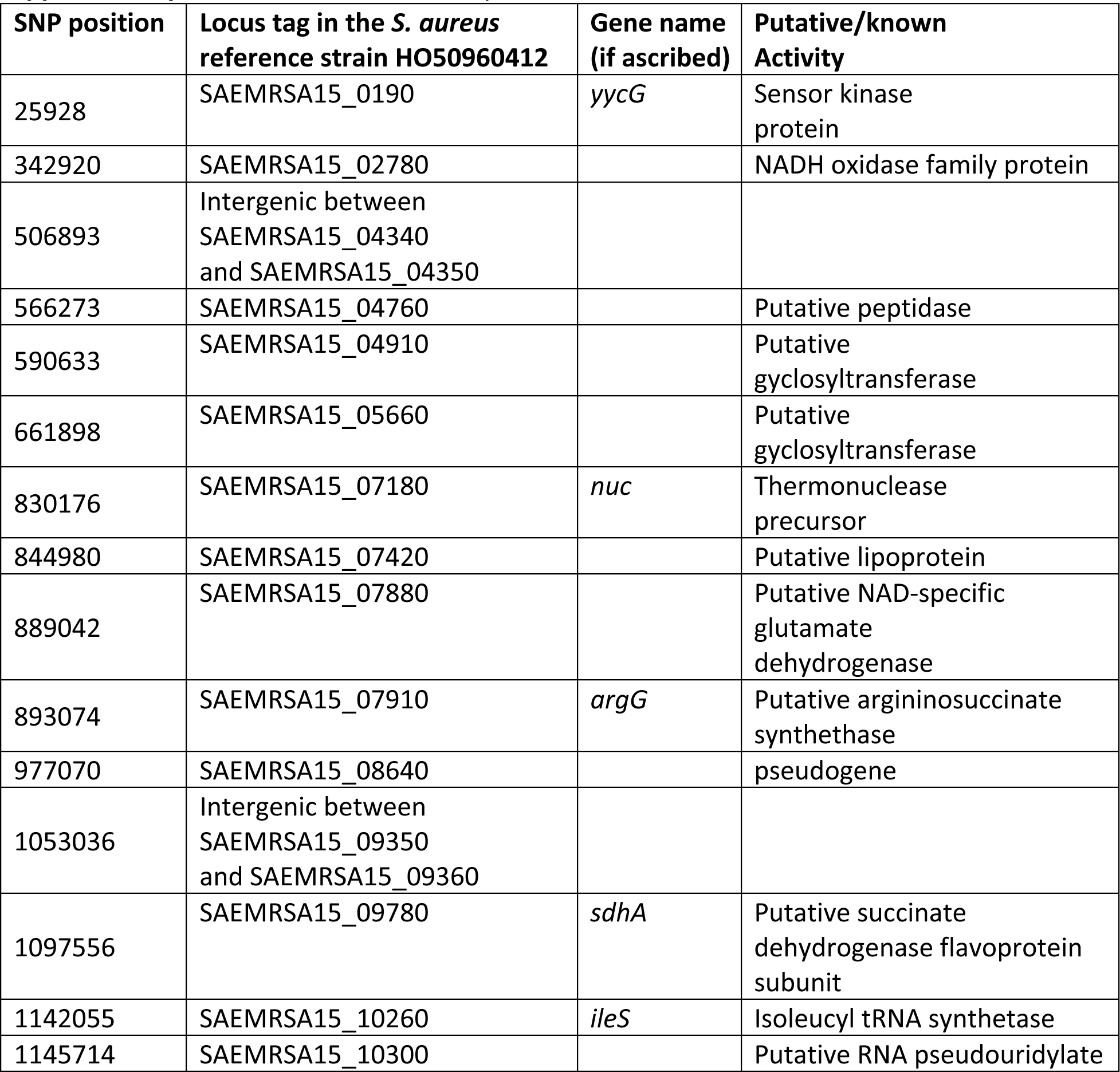
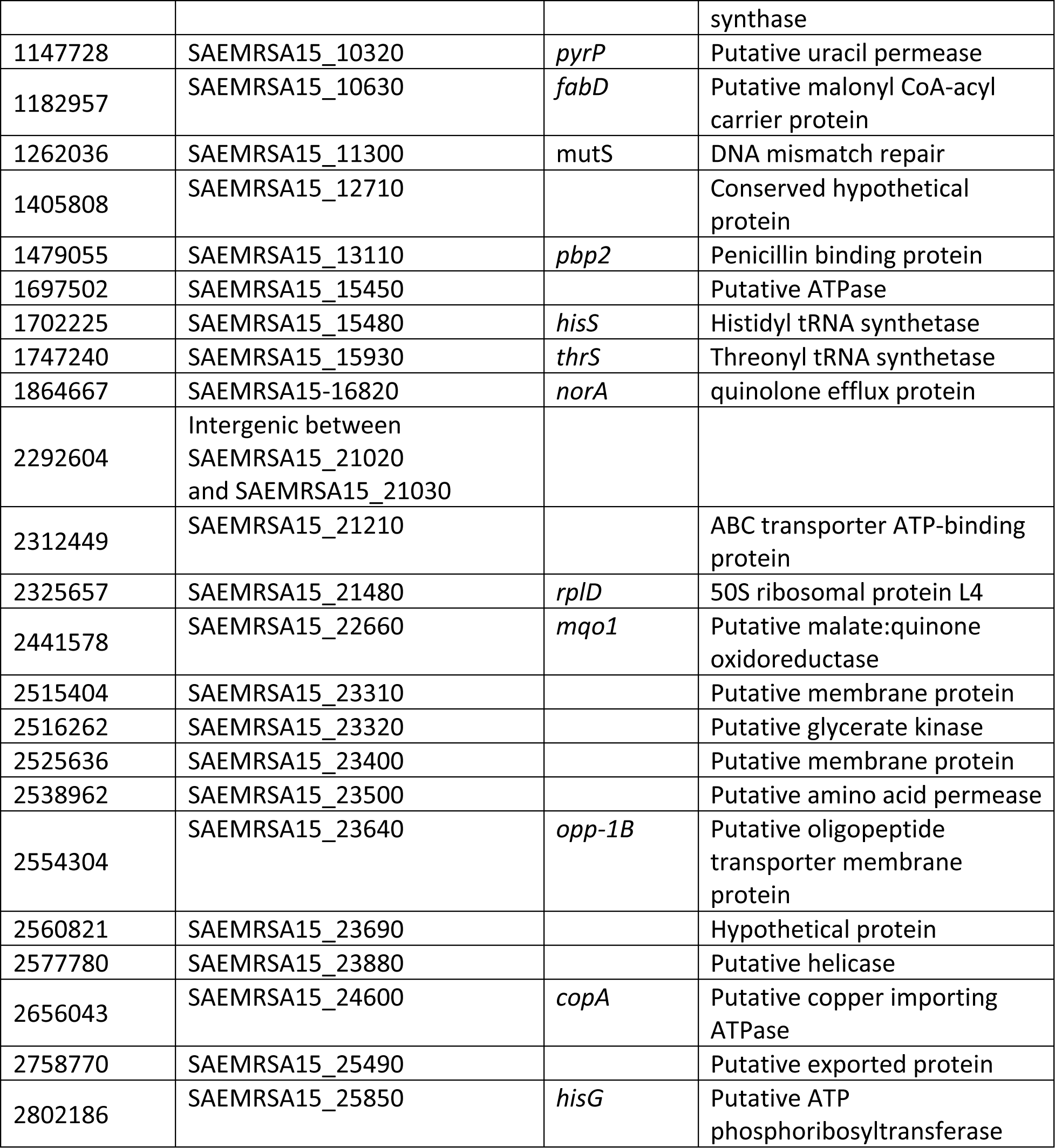
Loci associated by GWAS with biofilm formation.

**Supplementary Table 3:** Bacterial loci identified by the model as important in predicting SAB-4 associated mortality (fig. 4c).

| Allele | Locus tag in the *S. aureus* reference strain HO50960412 | Gene name (if ascribed) | Putative/known function |
| --- | --- | --- | --- |
| 398966 | SAEMRSA15_03350 |  | nitroreductase family protein |
| 142543 | SAEMRSA15_01150 | *capA* | capsular polysaccharide synthesis enzyme |
| 1691885 | intergenic between<br>SAEMRSA15_15380 and 15390 |  |  |
| 2013094 | SAEMRSA15_18250 |  | putative prephenate<br>dehydratase |
| 1335028 | SAEMRSA15_12030 | *grlA* | topoisomerase IV subunit A |
| 344051 | SAEMRSA15_02790 |  | putative bacterial luciferase<br>family protein |
| 254115 | SAEMRSA15_02010 |  | putative PTS transport system,<br>IIBC component |
| 1196127 | SAEMRSA15_10750 |  | putative membrane protein |
| 1348351 | SAEMRSA15_12130 |  | prephenate dehydrogenase |
| 1566333 | intergenic between<br>SAEMRSA15_14090 and 14100 |  |  |
| 72400 | SAEMRSA15_00580 |  | LysR-family regulatory protein |
| 1768164 | SAEMRSA15_16080 | *pfkA* | 6-phosphofructokinase |
| 2092793 | intergenic between<br>SAEMRSA15_19290 and 19300 |  |  |
| 165352 | SAEMRSA15_01380 |  | putative lipoprotein |
| 834688 | SAEMRSA15_07270 |  | phosphoglycerate mutase<br>family protein |

